# Exploring the evolution and adaptive role of mosaic aneuploidy in a clonal *Leishmania donovani* population using high throughput single cell genome sequencing

**DOI:** 10.1101/2020.03.05.976233

**Authors:** Gabriel H. Negreira, Pieter Monsieurs, Hideo Imamura, Ilse Maes, Nada Kuk, Akila Yagoubat, Frederik Van den Broeck, Yvon Sterkers, Jean-Claude Dujardin, Malgorzata A. Domagalska

## Abstract

Maintenance of stable ploidy over continuous mitotic events is a paradigm for most higher eukaryotes. Defects in chromosome segregation and/or replication can lead to aneuploidy, a condition often considered deleterious. However, in *Leishmania*, a Protozoan parasite, aneuploidy is a constitutive feature, where variations of somies represent a mechanism of gene expression adaptation, possibly impacting phenotypes. Strikingly, clonal *Leishmania* populations display cell-to-cell somy variation, a phenomenon named mosaic aneuploidy (MA). However, until recently, no method was available for the determination of the complete karyotype of single *Leishmania* parasites. To overcome this limitation, we used here for the first time a high-throughput single-cell genomic sequencing (SCGS) method to estimate individual karyotypes of 1560 promastigote cells in a clonal population of *Leishmania donovani*. We identified 128 different karyotypes, of which 4 were dominant. A network analysis revealed that most karyotypes are linked to each other by changes in copy number of a single chromosome and allowed us to propose a hypothesis of MA evolution. Moreover, aneuploidy patterns that were previously described by Bulk Genome Sequencing as emerging during first contact of promastigotes populations with different drugs are already pre-existing in single karyotypes in the SCGS data, suggesting a (pre-)adaptive role of MA. Additionally, the degree of somy variation was chromosome-specific. The SCGS also revealed a small fraction of cells where one or more chromosomes were nullisomic. Together, these results demonstrate the power of SCGS to resolve sub-clonal karyotype heterogeneity in *Leishmania* and pave the way for understanding the role of MA in these parasites’ adaptability.

**Update: 25^th^ May 2021:** A revision of the present preprint was released in BioRxiv on 11^th^ May 2021 (https://www.biorxiv.org/content/10.1101/2021.05.11.443577v2). In the new version, we included two extra samples in our single-cell genome sequencing (SCGS) analysis – the BPK081 cl8 clone (a nearly euploid strain), and a population consisting of a mixture of four *L. donovani* strains which was used as control for high levels of mosaicism in aneuploidy and for estimation of doublets. We also upgraded the bioinformatics pipeline to determine single-cell karyotypes and performed new fluorescence in situ hybridization (FISH) analysis. The new findings observed especially in the BPK081 cl8 led to a reformulation of the text, a new hypothesis for the evolution of mosaicism and a general restructuring of the article. Therefore, the present preprint is obsolete. Please refer to the new preprint entitled “High throughput single cell genome sequencing gives insights in the generation and evolution of mosaic aneuploidy in *Leishmania donovani*” for more information.

## Introduction

Historically, cell populations were analyzed in bulk assuming that this provides the representative information on the biology and behavior of these cells in a given experimental set up. The existence of heterogeneity between single cells, even in clonal populations, has been acknowledged but for long it was not further explored due to lack of suitable methodologies, and also because of the assumption that this cell-to-cell variation is random and has no biological significance. In the last years, advances in single cell imaging, molecular biology and systems biology enabled analysis and quantification of differences between single cells, defining new cell types and their origin, and finally allowing the demonstration of functional relevance of this variation^1^. This cell-to-cell variability of clonal populations is also of essential importance for unicellular organisms, like *Staphylococcus aureus*^2^, or *Sacharomyces cerevisiae*^3^. Similarly, for digenetic protozoan parasites, such as *Plasmodium spp*. or *Trypanosoma cruzi*, cellular mosaicism was also demonstrated to be crucial for drug resistance or tissue tropism during the vertebrate host infection^4^.

*Leishmania* sp. are digenetic unicellular protozoan parasites responsible for a spectrum of clinical forms of leishmaniasis worldwide and causing 0.7-1 million new cases per year^5^. As other trypanosomatids, *Leishmania* belongs to the supergroup Excavata, one of the earliest diverging branch in the Eukaryota domain^6^. Thus, several molecular mechanisms considered canonical for eukaryotes are different in these parasites, including the genomic organization in long polycistronic units, the near absence of transcription initiation regulation by RNA polymerase II promoters with gene expression regulation happening mostly through post-transcriptional mechanisms^7^, and its remarkable genomic plasticity^8^. The genome of *Leishmania* sp is ubiquitously aneuploid, and high levels of chromosomal somy variation, as well as local gene copy number variation are found between all *Leishmania* species^9,10^. Moreover, these variations are highly dynamic and change in response to new environment, such as drug pressure, vertebrate host or insect vector^11^. Changes in somy, together with episomal gene amplifications, and not variation in nucleotide sequence are the first genomic modifications observed at populational level during the course of experimental selection of drug resistance, suggesting that they are adaptive^12,13^. Importantly, somy alterations are reflected in the level of transcriptome^11^, and -for a same life stage-to a great degree of proteome derived from genes located in polysomic chromosomes^14^. These findings further support the notion that *Leishmania* exploits aneuploidy to alter gene dosage and to adapt to different environment.

Cell-to-cell variation in *Leishmania* has been so far reported in the form of mosaic aneuploidy (MA). This phenomenon was demonstrated in several *Leishmania* species by means of fluorescence in situ hybridization (FISH) performed on a few chromosomes, which showed at least two somy states among individual cells^15^. As a consequence, thousands of different karyotypes are expected to co-exist in a parasite population^16^. This heterogeneity of aneuploidy provides a huge potential to generate diversity in *Leishmania* from a single parental cell, both quantitatively through gene dosage, but also qualitatively through changes in heterozygosity^17^. Thus, it is hypothesized that mosaic aneuploidy constitutes a unique source of adaptability to new environment for the whole population of parasites^16^.

Although MA has been demonstrated by FISH, its extent to all over the 35-36 chromosomes of *Leishmania*, its dynamics in constant as well as new environment and its potential role in adaptation to different environment remains to be determined. Accordingly, pioneer FISH-based studies should be complemented and refined by single cell genome sequencing (SCGS). In a previous study^18^, we made a first step in that direction by combining FACS-based sorting of single *Leishmania* cells with whole genome amplification (WGA) and whole genomic sequencing (WGS). In this pilot study, we evaluated different WGA and bioinformatic methods and detected 3 different karyotypes among 28 single cells of *L. braziliensis*^18^. Here, we made one step beyond, applying a droplet-based platform for single cell genomics, in order to undertake the first high-throughput study of MA in *Leishmania*.

## Material and Methods

### Parasites

*L. donovani* promastigotes of the strain MHOM/NP/03/BPK282/0 clone 4 (further called BPK282, reference genome of *L. donovani*) were maintained at 26°C on HOMEM medium (Gibco, ThermoFisher) supplemented with 20% Fetal Bovine Serum, with regular passages done every 7 days. The strain was submitted to SCGS 21 passages after cell cloning and analyzed by FISH 23 passages later.

### Cell Suspension Preparation for SCGS

BPK282 promastigotes at early stationary phase (day 5) were harvested by centrifugation at 1000rcf for 5 minutes, washed twice with PBS1X (without calcium and magnesium) + 0,04% BSA, diluted to 5×10^6^ parasites/mL and passed through a 5μm strainer to remove clumps of cells. After straining, volume was adjusted with PBS1X + 0,04% BSA to achieve a final concentration of 3×10^6^ parasites/mL. The absence of remaining cell duplets or clumps in the cell suspension was confirmed by microscopy.

### Single Cell partitioning, barcoding, WGA and sequencing

We used 4.2μL of the single-cell suspension as input to the 10X Chromium™ Single Cell CNV Solution protocol (10X Genomics), targeting a total of 2000 sequenced cells according to the manufacturer’s instructions. Individual cells were portioned and incapsulated in a hydrogel matrix using a microfluidic chip and the Chromium Controller (10X Genomics). During cell partitioning, each cell was combined with the Cell Bead (CB) Polymer™ and the Cell Matrix™ reagents, forming the CBs. CBs were then incubated overnight at 21°C and 1000RPM on a thermomixer for homogeneous hardening of the CB polymer. The individually incapsulated cells were lysed, releasing the genomic DNA (gDNA) inside the CB. In a second microfluidic chip, each CB was individually joined with a Gel Bead (GB) containing multiple copies of one of the ~750.000 unique 10X barcodes™, together with an enzyme mix used in downstream Whole Genome Amplification (WGA) and ligation of 10X barcodes. The CB-GB joining was performed with a partitioning oil, forming an emulsion where each droplet (GEMs) contain a single CB linked to a unique GB. Inside the GEMs, the CB and GB were disrupted, and isothermal WGA with random hexamers followed by ligation of the 10X barcode to the amplified gDNA molecules was carried out. Then, GEMs were disrupted, amplified DNA molecules were pooled and processed for Illumina sequencing with the addition of P5 and P7 adaptors and a sample index. Sequencing of the library was performed at Genomics Core Leuven (Belgium), with a NextSeq High Output kit (Illumina) platform with 2 x 150 read length.

### CNV Calling and Somy Estimation

Reads were associated to each sequenced cell based on their 10X-barcode sequence and mapped to a customized version of the reference *L. donovani* genome LdBPKv2^11^ (available at ftp://ftp.sanger.ac.uk/pub/project/pathogens/Leishmania/donovani/LdBPKPAC2016beta/), where Ns were added to the ends of chromosomes 1 to 5 to reach the 500kb minimum size allowed by the Cell Ranger DNA^™^ pipeline (10X Genomics). The normalized read depth (NRD) within adjacent, non-overlapping 80kb bins was used to calculate copy number values (referred here as CNV values) in 80kb intervals, using the version 1.1 of the Cell Ranger DNA^™^ pipeline^19^. CNV values represent the integer NRD of each bin after scaling of NRD values by the baseline ploidy factor (S factor) of each cell determined by the scaling algorithm of the pipeline^20^. We used the arguments min-soft-avg-ploidy and max-soft-avg-ploidy set to 1.9 and 2.1, respectively, to encourage the scaling algorithm to establish cells baseline ploidy to 2. This was done to avoid overscaling artifacts generated by the scaling algorithm that were observed in some cells when these options were not used. CNV values were visualized in the Loupe scDNA Browser (10X Genomics)^21^, with cells arranged in 512 clusters, the maximum number of clusters allowed by the software. These clusters were composed of 1 to 7 cells each that were grouped together based on CNV values similarities^22^. CNV values of each 80kb bin of all 512 clusters were exported as a csv file and the average CNV values of intrachromosomic bins were used to estimate chromosomal somy. Bins mapping to high copy number loci such as the H-locus (chromosome 23) and M-locus (chromosome 36)^23^ were excluded from the calculation. Bin-to-bin variability was estimated by the normalized standard deviation of CNV values of intrachromosomal bins. Each cell cluster received a bin variability score based on the average bin-to-bin variability of all chromosomes. Clusters displaying a bin variability score higher than 0.05 were excluded. Remaining clusters were ungrouped, with somy values of each individual cell corresponding to the somy values estimated for their original clusters.

### Karyotypes identification and network analysis

In order to identify different karyotypes present in the sequenced population, somy values of each cell were rounded to their closest integer. Cells displaying the same rounded somy values for all chromosomes were considered as having the same karyotype. Each unique karyotype identified in the population received an identifier composed by the concatenated rounded somy values of all chromosomes, from chromosome 1 to chromosome 36 (e.g., “222232223222222322222232232222423232”). Unique karyotypes were numerically named according to their frequency in the sequenced population.

To perform a network analysis, a pairwise distance matrix was built based on the number of different chromosomes between all karyotype and used to generate a randomized minimum spanning tree with 100 randomizations, using the Pegas R package^24,25^.

### DNA probes and fluorescence in situ hybridization (FISH)

DNA probes were either cosmid (L549 specific of chromosome 1) or BAC (LB00822 and LB00273 for chromosomes 5 and 22 respectively) clones that were kindly provided by Peter Myler (Seattle Biomedical Research Institute) and Christiane Hertz-Fowler (Sanger Centre). DNA was prepared using Qiagen Large-Construct Kit and labelled with tetramethyl-rhodamine-5-dUTP (Roche Applied Sciences) by using the Nick Translation Mix (Roche Applied Sciences) according to manufacturer instructions. Cells were fixed in 4% paraformaldehyde then air-dried on microscope immunofluorescence slides, dehydrated in serial ethanol baths (50–100%) and incubated in NP40 0,1 % for 5 min at RT. Around 100 ng of labelled DNA probe was diluted in hybridization solution containing 50% formamide, 10% dextran sulfate, 2× SSPE, 250 μg.mL^-1^salmon sperm DNA. Slides were hybridized with a heat-denatured DNA probe under a sealed rubber frame at 94°C for 2 min and then overnight at 37°C and sequentially washed in 50% formamide/2× SSC at 37°C for 30 min, 2× SSC at 50°C for 10 min, 2× SSC at 60°C for 10 min, 4× SSC at room temperature. Finally, slides were mounted in Vectashield (Vector Laboratories) with DAPI. Fluorescence was visualized using appropriate filters on a Zeiss Axioplan 2 microscope with a 100× objective. Digital images were captured using a Photometrics CoolSnap CCD camera (Roper Scientific) and processed with MetaView (Universal Imaging). Z-Stack image acquisitions (15 planes of 0.25μm) were systematically performed for each cell analyzed using a Piezo controller, allowing to view the nucleus in all planes and to count the total number of labelled chromosomes. Around 200 cells [187-228] were analyzed per chromosome.

## Results

In this study, we submitted the clonal population of BPK282 promastigotes to a high throughput, droplet-based SCGS method (Chromium™ CNV Solution - 10X Genomics). In total, 148,7M reads were generated, of which 49,7M were mapped to 1703 cells after removal of duplicated reads. The average effective coverage depth per cell was 0,14x, with only 2 cells displaying a depth coverage higher than 1x (**Figure S1A)**. However, read mapping was evenly distributed across chromosomes in general(**Figure S1B**), which allows somy estimation despite low depth^18^. From 512 clusters, 452 passed the bin-to-bin variability filtering, representing a total of 1560 cells. Most eliminated clusters displayed an average somy close to 1 (**Figure S1C**).

### Copy number variation of individual chromosomes

The somies of the 36 *L. donovani* chromosomes in the 1560 filtered cells are depicted in **Figure 1A**. Of these 36 chromosomes, 7 were predominantly trisomic (chromosomes 5, 9, 16, 23, 26, 33 and 35), while chromosome 31, the only chromosome that is extensively reported as polysomic (often tetrasomic) in all *Leishmania* species studied so far^9,26^, was also tetrasomic in most cells in our SCGS data. The other chromosomes were mostly disomic. These observations are very similar to the populational aneuploidy pattern estimated by BGS of the BPK282 reference strain (**Figure 1A, top heatmap, and Figure S2**). Notably, chromosome 13, which display an intermediate somy value in the Bulk Genomic Sequence (BGS) data, was found as disomic and trisomic at high proportions in the SCGS. Moreover, our SCGS data revealed that high cell-to-cell somy variation was chromosome-specic. For instance, chromosomes 21 and 18 were disomic in 99,23% and 99,42% of the cells respectively, while chromosomes 13, and 35 were the most variable ones, with 33,54% and 25,24% of the cells displaying a copy number divergent from the most frequent somy.

**Figure 1.**
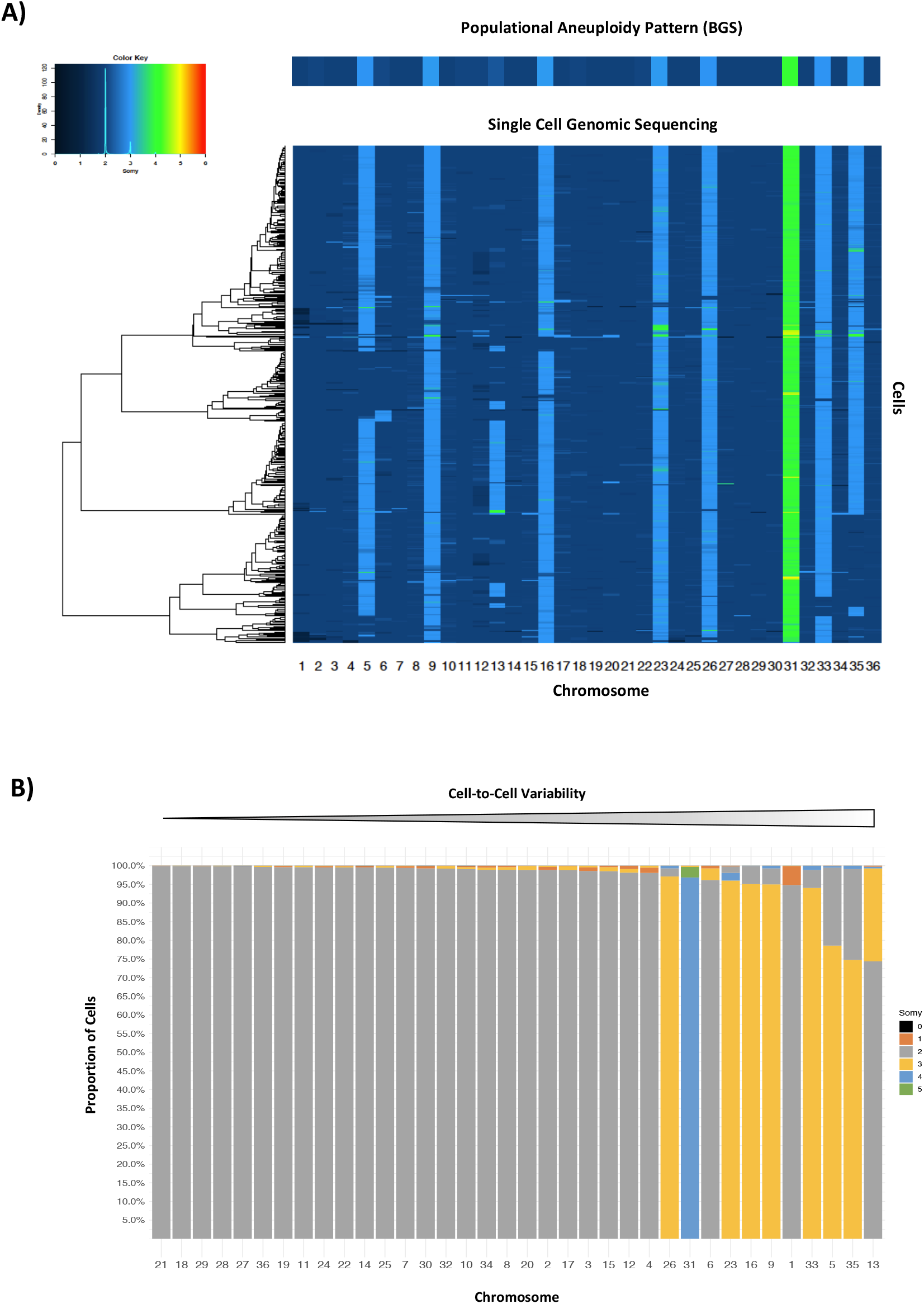
**A)** Somy values of all 36 chromosomes of all 1560 cells in the BPK282 clonal population. The color key inset displays the distribution of somy values in the SCGS data. A heatmap based on BGS data of BPK282 is displayed on the top. **B)** Proportion of somies found for each chromosome. Chromosomes are arranged, from left to right, by cell-to-cell variability, defined as the proportion of cells displaying a somy different from the most frequent one. For each column, somy bars are stacked by the frequency they happen in the population.

In the filtered data, we identified a cell where 4 chromosomes were estimated as nullisomic (**Supplementary table 1 - kar89)**. The bam file of this cell shows that these chromosomes had no sequenced reads mapping to them (**Figure 2A**, first plot), indicating that they were absent in this cell. An investigation of the bam files of all cells, including the ones eliminated after data filtering, revealed a total of 12 cells displaying at least one chromosome with no reads mapping to it, of which 11 displayed a high intrachromosomal bin-to-bin variability of the mapped chromosomes and were excluded during data filtering.

**Figure 2.**
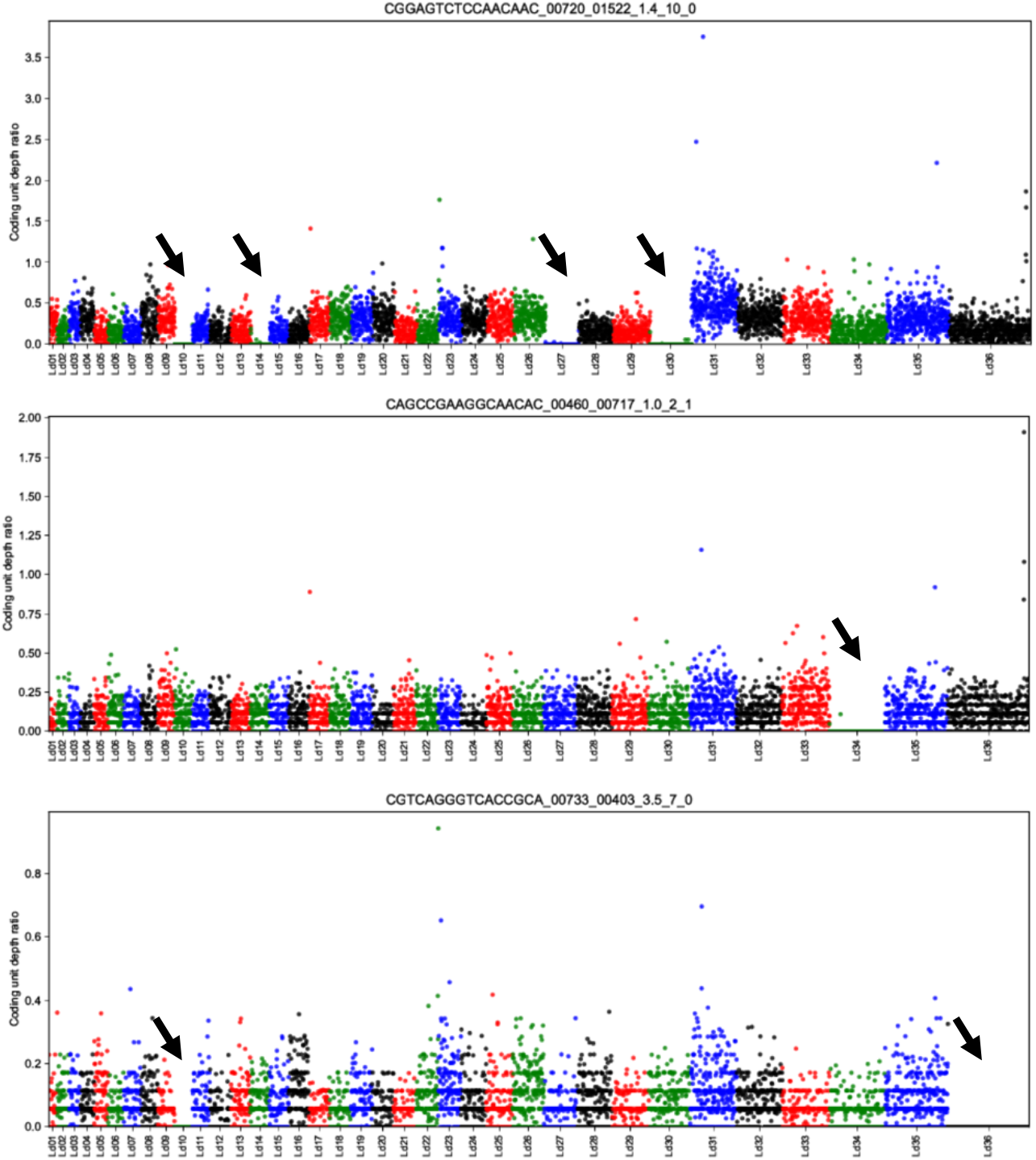
Three examples of individual cells, identified by their 10X barcode sequence (top of each plot), where one or more chromosome is nullisomic. Manhattan plot displays average depth per 5kb. Nullisomic chromosomes are indicated with a black arrow. When present, single reads mapping to nullisomic chromosomes are reads composed by repetitive nucleotides that can be mapped to multiple regions of the genome.

### Karyotype mosaicism

A karyotype was defined as any unique combination of rounded somy values for all chromosomes identified in the sequenced cells after data filtering. According to this definition, 128 different karyotypes were identified amid 1560 cells. The most frequent karyotype (kar1) was found in 431 cells, representing 27,65% of the sequenced population; this karyotype is equivalent to the average karyotype observed by BGS of BPK282. Four karyotypes are dominant in the population, representing 62,5% of cells (**Figure 3A**). Noteworthy, the most dominant karyotypes displayed little somy difference between each other. For instance, kar2, 3 and 4 differ from kar1 by changes in somy in a single chromosome, i.e. chromosomes 13, 35 and 5 respectively, while kar5 diverge from kar2 by a disomy in chromosome 35 (**Figure 3B**). A network analysis showed that most karyotypes diverge from another by a difference in a single chromosome, displaying an interesting pattern where most karyotypes can be linked back to kar1 by single chromosome-change steps (**Figure 4**, black pies). Noteworthy, kar1 also displays the biggest number of secondary karyotypes directly linked to it by somy changes in single chromosomes, including the other 3 dominant karyotypes. Karyotypes which the closest relatives were at two or more chromosome changes apart (red pies) were only present at frequencies lower than 0,44%. Moreover, the only karyotype with nullisomic chromosomes that passed the data filtering criteria (kar89) display several chromosomes with somies different from most karyotypes, with the closest one (kar32) being 18 chromosomes somy changes apart from it.

**Figure 3.**
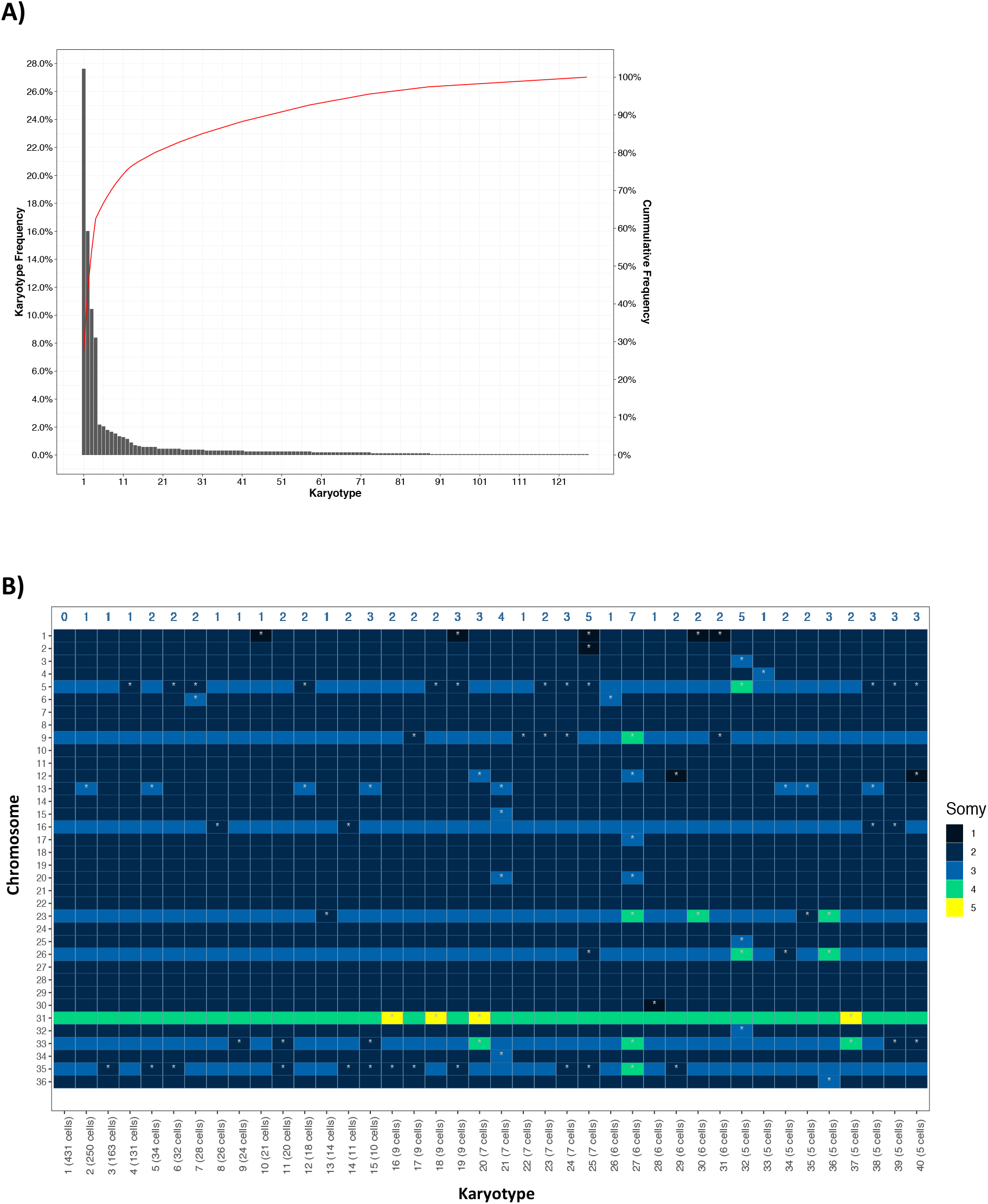
Distribution and profile of different karyotypes identified among 1560 BPK282/0 cl4 promastigotes. **A)** Frequency distribution of **a**ll the 128 different karyotypes identified in the SCGS data. Red line indicates the cumulative frequency in the secondary axis. **B)** Heatmap displaying the somy values off each chromosome in the 40 most frequent karyotypes. Karyotypes are ordered from left to right and numbered according to their frequency in the population. Chromosomes in which somies diverge from the most frequent karyotype (Karyotype 1) are marked with a white asterisk. Blue numbers in the top part of the graph indicate the number of chromosomes with somy different from the first karyotype.

**Figure 4.**
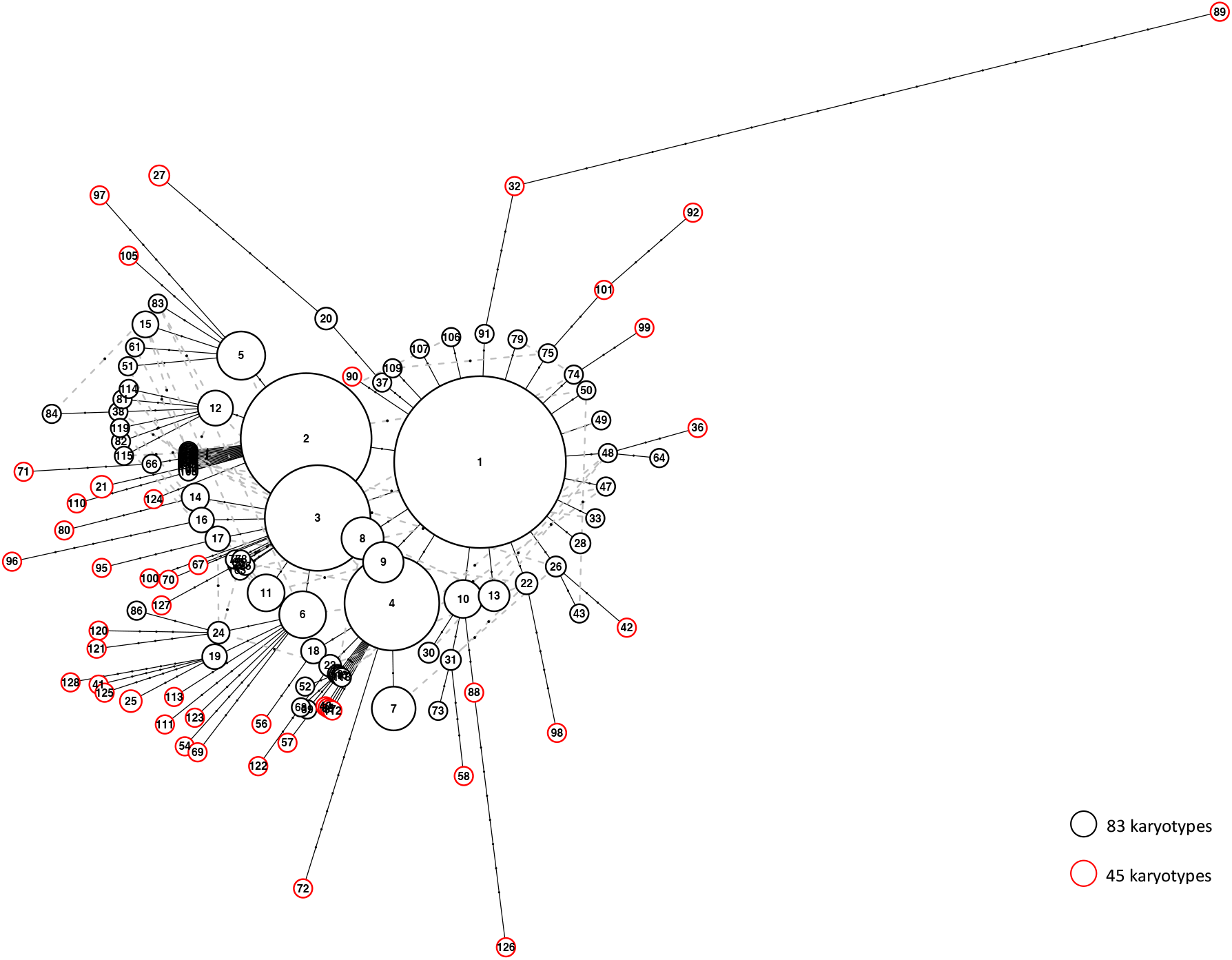
Haplotype network between all 128 karyotypes. The dots in the link lines represent the number of different chromosomes between 2 karyotypes. Dashed grey lines indicate alternative links between karyotypes that diverge by a single chromosome (black pies). Red pies highlights karyotypes where the closest related karyotype display different somy in at least 2 chromosomes. The size of each pie is proportional to the number of cells displaying a given karyotype. The smallest pies represent a total of 7 or less cells.

### Pre-existing karyotypes selected during early environmental adaptation

Previous studies have demonstrated through BGS that *in vitro Leishmania* populations display changes in average aneuploidy patterns of the cell population as one of the first genomic alterations when submitted to environmental changes, including hosts infection and drug pressure^11–13,27^. However, it is still unknown if these variations in somies observed at populational level are generated by selection of karyotypes pre-existing in subpopulations or represent *de novo* alterations induced by the new environmental pressure. To address this issue, we revisited previously published BGS data where the same BPK282 clonal promastigote population was used as model in drug selection experiments. We evaluated in our SCGS data if changes in aneuploidy patterns that emerge during adaptation to different drug pressures at populational level are already present in single karyotypes in the BPK282 population.

In the 3 revisited drug resistance selection studies, the same pattern was observed. First rounds of selection led to changes in the average populational aneuploidy profile that were equivalent to single karyotypes present in our SCGS data (**Figure 5**). Authors^13^ reported a reduction in the somy of chromosomes 13 and 35 in a BPK282 population exposed to 3μM and 6μM of Miltefosine. The observed populational aneuploidy pattern is similar to kar9, found in 24 cells in our SCGS data, where chromosomes 13 and 35 are disomic (**Figure 5A**). A similar observation happened when BPK282 promastigotes were exposed to 2μM and 4μM of Paramomycin in another study^27^, where the somy changes are also similar to kar9 (**Figure 5B**). Finally, *in vitro* SbIII selection experiments with BPK282 led to changes in populational average somies which were similar to karyotypes 3, 4, or 6 among 3 replicates exposed to 48μM of the drug ^12^ (**Figure 5C**). However, in these 3 studies, higher drug concentrations led to changes in the average populational somies that are not represented by single karyotypes in our data.

**Figure 5.**
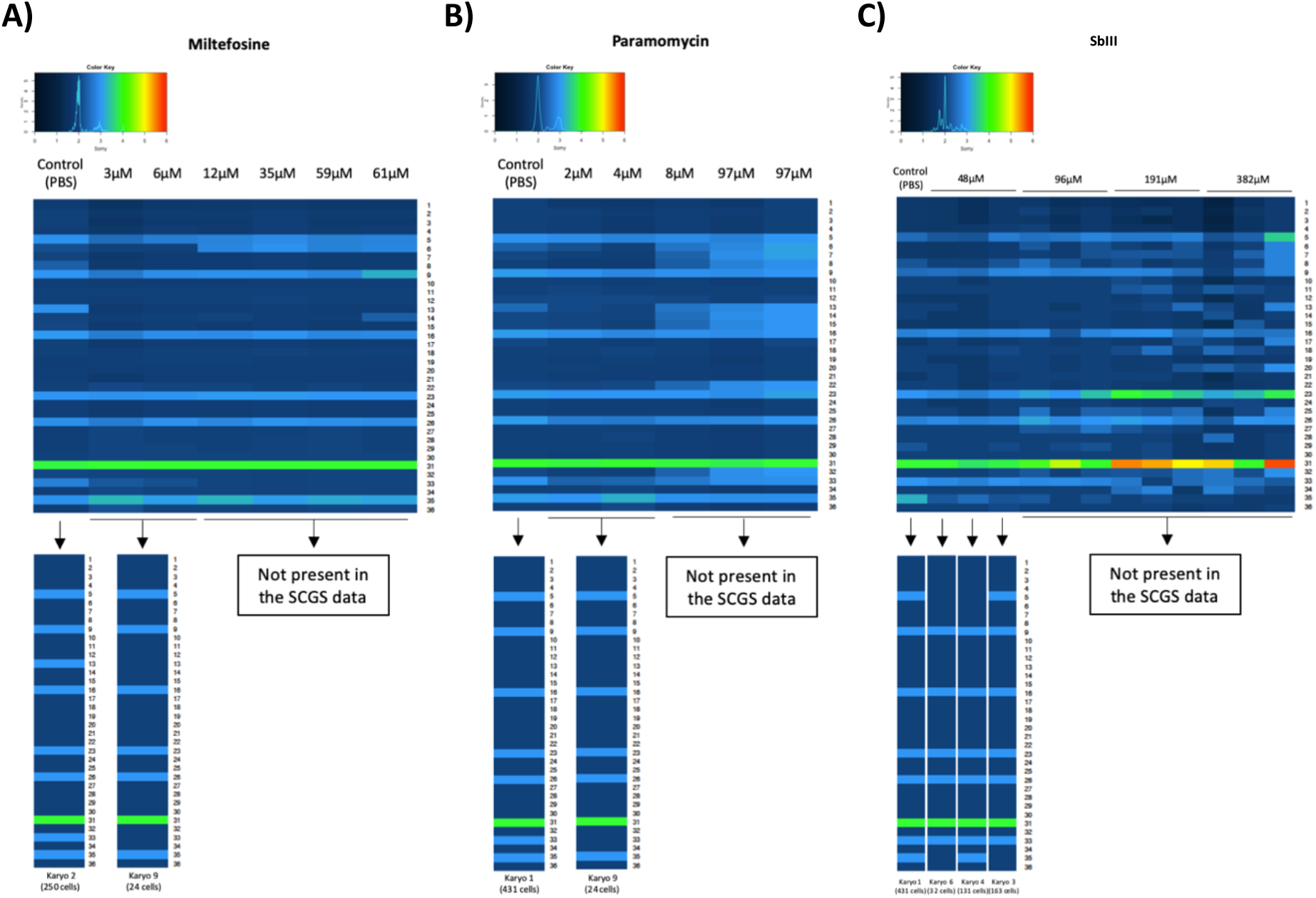
Comparison between previously published somy estimation by BGS of BPK282/0cl4 strain during early adaptation to drug pressure (upper heatmaps) and individual karyotypes found in the SCGS data (bottom heatmaps). BGS somy values were published elsewhere for Miltefosine^13^, Paramomycin^26^ and SbIII^12^.

### Validation of the SCGS results with FISH

As a validation to our SCGS data, we also evaluated the somy of 3 chromosomes in 187-228 BPK282 cells using FISH. We observed very similar somies distributions in both methods (**Figure 6**). For instance, chromosome 5, the most variable of the 3 assessed chromosomes, was estimated as trisomic in 78,5% of the cells in the SCGS data, and 74,5% by FISH, while it was disomic in 20,9% in SCGS and 22,6% in FISH. Chromosome 1 was estimated as disomic in 94,8% of the cells in the SCGS and 96,2% in FISH, with a smaller fraction of cells being monosomic (5,1% in SCGS and 3,2% in FISH). Chromosome 22 had slightly more divergent values between both methods, being disomic in 99,5% in SCGS and 92,1% in FISH, with a fraction of 7,9% of cells that were found as monosomic in FISH while monosomy for chromosome 21 was found in only 0,19% of the cells in SCGS. In general, the similarities between the distributions found in both methods supported the reliability of SCGS.

**Figure 6.**
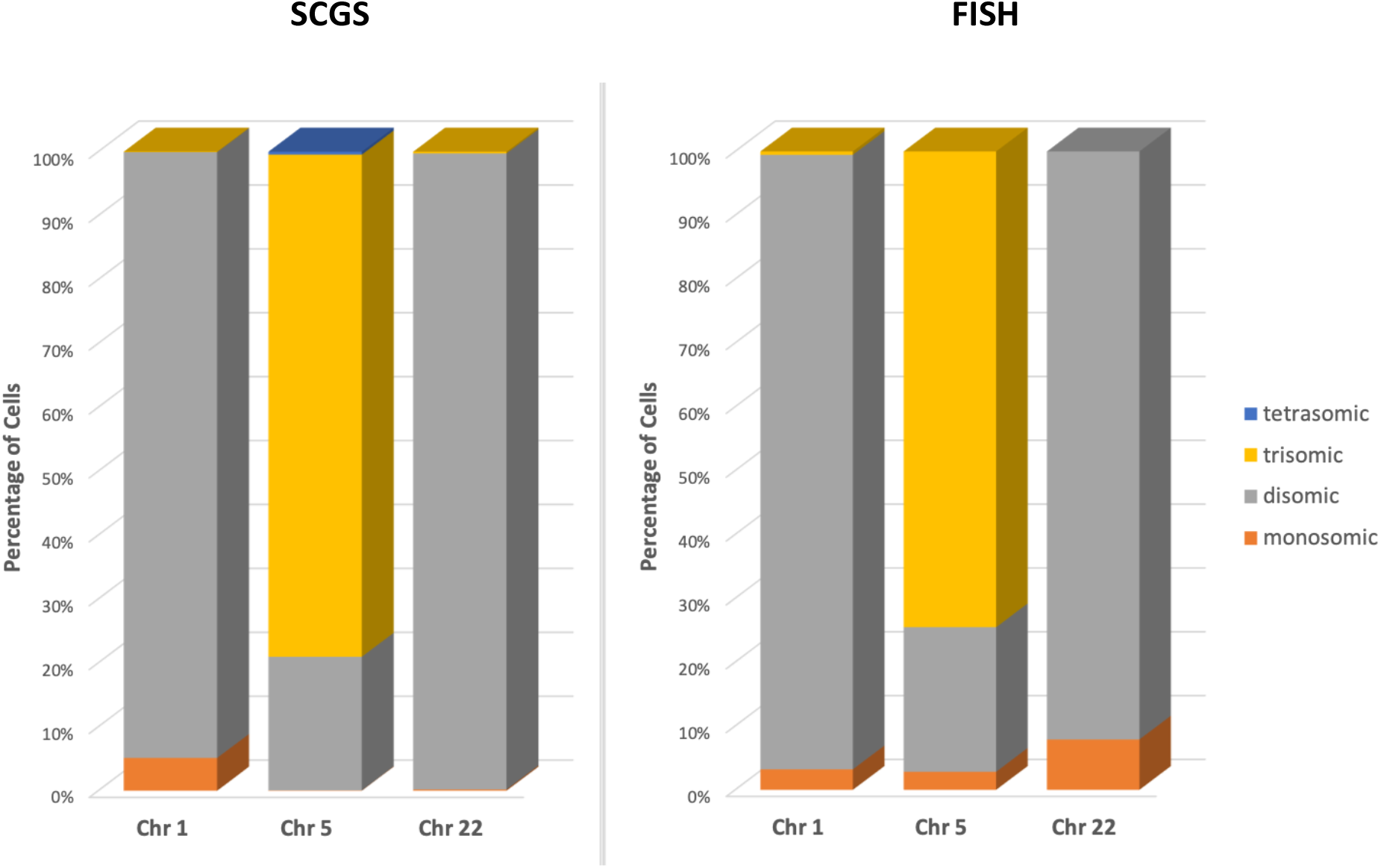
Validation of the SCGS data using FISH. The proportions of cells displaying monosomic, disomic, trisomic and tetrasomic chromosomes 1, 5 and 22 were evaluated by FISH and compared to the same proportions found in the SCGS data.

## Discussion

The parallel whole genomic sequencing of 1560 *Leishmania* promastigotes reported here allowed us to reveal for the first time a complex mosaicism of complete karyotypes in *Leishmania* with unprecedent resolution. By estimating the somy of all 36 chromosomes in each sequenced single parasite, we could determine which chromosomes were more prone to cell-to-cell somy variability, which were the frequencies of each identified karyotype in the clonal population and the divergencies between these karyotypes. Our data was further supported by FISH, which is a well establish method for the quantification of individual chromosome copy number in single *Leishmania* cells^28^. Thus, this work represents the first high throughput SCGS study of MA in *Leishmania*.

### Performance and challenges of the SCGS method

In general, the whole genome amplification (WGA) method used in the Chromium™ Single Cell CNV solution seems to generate an even genome coverage, which has been demonstrated as a more relevant aspect than coverage depth for accurate somy estimation^18,29^. However, a low percentage of cells displaying a higher read count variability might reflect on inaccurate CNV values, and therefore, unreliable karyotype determination. The Cell Ranger DNA™ pipeline uses an hierarchical clustering algorithm to combine the read counts of cells with similar CNV values to enhance the resolution and reliability of CNV values determination^22^, thus greatly reducing the number of faulty karyotypes. Yet, cells with slightly different karyotypes might be clustered together due to inaccurate single-cell CNV values estimation, leading to a cluster somy pattern that does not reflect a true biological karyotype. Since we noticed that artificial karyotypes were caused by chromosomes with high variation in intrachromosomal CNV values, we applied an addition data filtering step where we eliminated clusters displaying a high intrachromosomic bin-to-bin variability. This led to the removal of 56 clusters (143 cells), corresponding to 51 karyotypes. Most of the eliminated karyotype displayed somy patterns that were highly distinct from the ones found in the remaining karyotypes after data filtering, probably due to inaccurate somy estimation caused by the high intrachromosomic bin-to-bin variability.

Most eliminated clusters had the majority of CNV values assigned to 1. Since CNVs are calculated by the relative read depth of the bins, the Cell Ranger DNA™ pipeline uses an algorithm that determines the baseline ploidy of each cell by scaling the normalized read depth values of each bin by a factor S that satisfies the condition where all normalized read depth values are integers multiple of S^20^. Candidate values for S are heuristically determined following a rule where if the cell displays the same copy number in most of the genome, S is set to a value that leads to an average ploidy close to 2, otherwise, in the presence of high variable CNV values, S is determined as the lowest value that satisfies the condition mentioned above. Thus, it is likely that the high variation of local CNV values present in these clusters, reflecting in a high intrachromosomic bin-to-bin variability, led to an unprecise determination of the S factor by the scaling algorithm, leading to an average ploidy in these clusters close 1. Indeed, the software cannot identify haploid cells since it is based on a normalized read depth quantification method. Haploid cells could be identified by the evaluation of heterozygous SNPs in single cells. However, the read depth per cell of our data does not allow such analysis.

The Cell Ranger DNA^™^ pipeline was developed to handle mammal genomes^30^. The nuclear genome of *Leishmania* is about 100 times smaller than human’s, and even the biggest *L. donovani* chromosome (Chr36, 2,768Mbp^31^) is still 17 times smaller than the smallest human chromosome (Chr21, 46,7Mbp). With the 80kb bin size determined by the pipeline in this data, small chromosomes such as Chr1, Chr2 and Chr3 are represented by 3 to 5 bins. However, this does not seem to introduce a variability bias to small chromosomes in general, since the observed cell-to-cell somy variability does not correlate to chromosome size. For instance, Chr4, Chr5 and Chr6 are all represented by 6 bins, and while Chr5 was found as triploid and diploid in high proportions for both states, Chr4 and Chr6 were among the less variable chromosomes (**Figure 1B**). Conversely, Chr34 and Chr35 had 24 and 27 bins respectively, and while the former was found as disomic in more than 95% of the cells, the later was estimated as disomic in 26,9% and trisomic in 71,4% of the cells. There is the possibility, however, that the very low number of bins for chromosome 1 (3 bins) introduced a bias towards monosomy.

On the other hand, the reduced size of *Leishmania* genome means that higher coverage depth per cell are achieved with relatively lower total sequence depth. The average 29.191 mapped deduplicated reads per cell resulted in an average coverage depth/cell of 0,14x. For human cells, the recommended 750.000 reads/cell is expected to yield a coverage depth of ~0,05x/cell, allowing CNVs assessment at 2mb resolution only^32^. This means that for each bin, a higher number of reads are expected in *Leishmania*, increasing CNV estimation resolution. Therefore, the higher coverage depth/cell seem to compensate the lower number of bins per chromosome in *Leishmania* genome.

### SCGS data supports errors in chromosome replication as main drivers of mosaicism

In a previous FISH-based study, authors reported asymmetrical chromosome allotments (ACA) in dividing nuclei as the cause of MA^15^. Remarkably, in all cells displaying ACA, the total copy number of the evaluated chromosome in the daughter cells were always odd (“3+2” or “2+1”). This observation suggests that mosaicism in aneuploidy is generated by defects in replication of chromosomes during cell division, as discussed by the authors, and not by errors in chromosome segregation, where the expected number of copies between daughter cells would be even. In our SCGS data, a pairwise comparison between all karyotypes that diverged from another by a single chromosome revealed that complementary karyotypes that could represent mis-segregation event are very rare. Of 127 possible pairs, only 7 pairs could be explained by a mis-segregation event (**Figure 7 and Supplementary table 2**). The other 120 pairs displayed a “2+1”, “2+3”, “3+4” or “4+5” pattern in the divergent chromosome, supporting the hypothesis that such karyotypes were probably generated by under-or over-replication of single chromosomes during mitosis.

**Figure 7.**
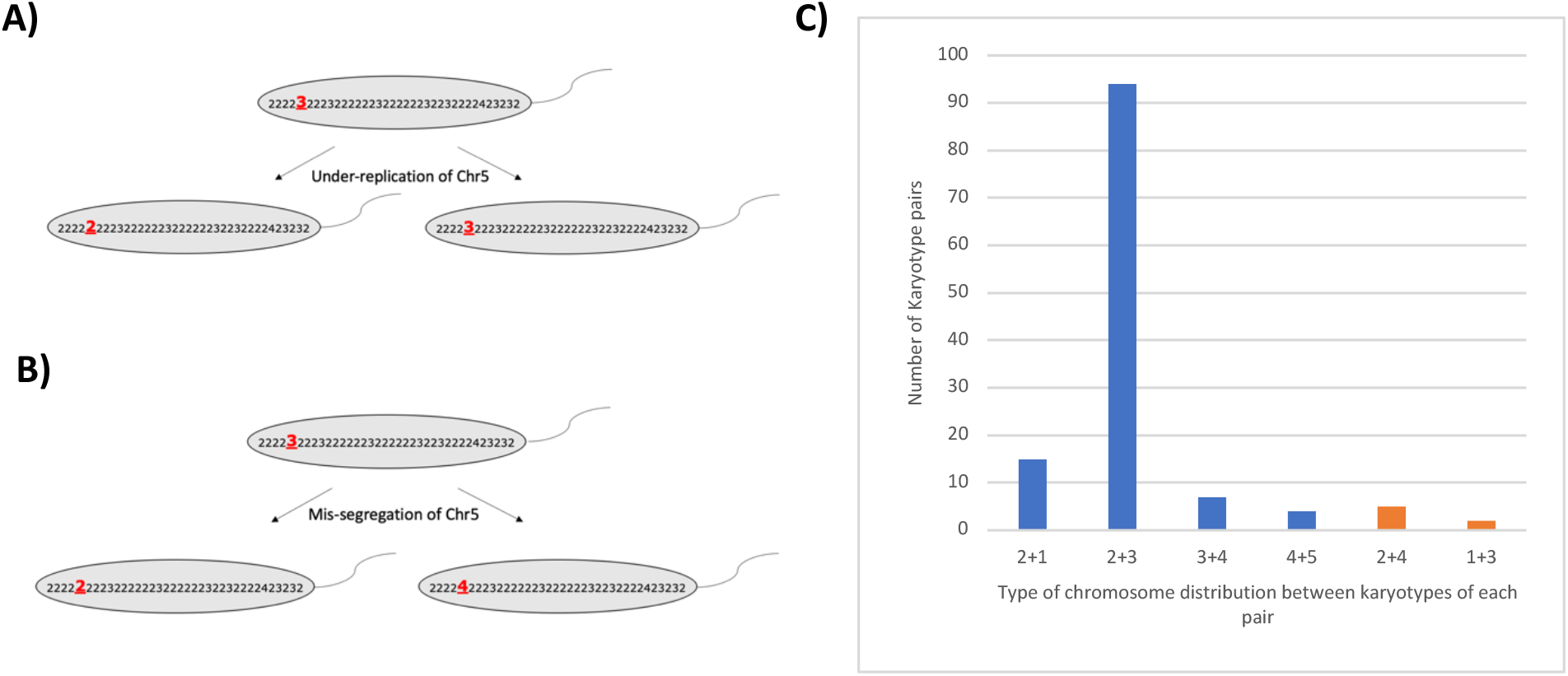
Karyotypes found in the SCGS data support errors in chromosome replication during mitosis as main drivers of aneuploidy mosaicism. **A)** An example of an under-replication event that leads to two karyotype pairs displaying a “2+3” (odd) somy distribution in chromosome 5. **B)** A mis-segregation event in chromosome 5 would generate two karyotypes that display a “2+4” (even) somy distribution in the daughter cells. **C)** Karyotypes diverging by a single chromosome from the SCGS data were compared in pairs. The pattern of somies of the divergent chromosome in each pair is represented in the x axis. Odd patterns, representing putative under or overreplication events, are depicted in blue, while even patterns, a putative indication of missegregation events, are represented in orange.

One of the effects of ACA events is the possibility of some cells losing all copies of a chromosome (nullisomy). Among all 1703 cells sequenced, including the ones that were removed for further analysis, 12 had one or more chromosome which was nullisomic. This represents ~0,7% of the total, which is a relatively high frequency considering that an *in vitro Leishmania* culture can consist of millions of cells. The fact that almost all cells with an absent chromosome displayed a karyotype that was not similar to any other in the sequenced population (**Figure S4**) is possibly due to the high bin-to-bin variability found in these cells, which hampers CNV estimation. The noisy coverage of these cells could be an indicative of DNA degradation after cell death, potentially caused by the lack of one or more chromosomes, suggesting that nullisomy is lethal for these parasites. In this case, the relatively high rate of cells with absent chromosomes is an indicative that ACA events that lead to nullisomy happen at high frequencies. Indeed, the FISH analysis of dividing nuclei found that for 2 chromosomes (chromosomes 2 and 22), around 1% of the evaluated parasites were displaying a “1+0” distribution of chromosomes between sister nuclei in *L. major*^15^.

### Chromosome Somy Variability

Although pioneer FISH studies demonstrated that somy mosaicism was more prominent in some chromosomes than others, the complete somy landscape of all chromosomes in a *Leishmania* population was still unknown at single-cell level, since only 11 chromosomes were evaluated by FISH hitherto. Our SCGS data revealed that some chromosomes display a remarkable lack of cell-to-cell variability in the sequenced population, while others are more prone to mosaicism. Several of the less variable chromosomes also display low inter-strain variation between *L. donovani* isolates. For instance, chromosomes 10, 17, 18, 19, 21, 24, 25, 27, 28, 30, 34 and 36 were previously reported as disomic by BGS in more than 95% of 204 isolates in a previous study^33^, and were disomic in more than 98% of the cells in our SCGS data, suggesting a pressure to maintain these chromosomes as disomic, at least under standard *in vitro* conditions. Conversely, chromosomes 5, 13, 33 and 35, the most variable in the SCGS, are also present as disomic or trisomic at high proportions between these different isolates. This indicates that, in a given environment, somy variability is restricted to a specific group of chromosomes. One possibility is that gene contents of some chromosomes are more tolerant to dosage imbalance than others, suggesting that selective pressure maintain somy stability in some chromosomes while allow more plasticity in others. Moreover, the WGS of 204 *L. donovani* isolates revealed that only chromosome 34 was consistently disomic in all strains^33^. However, BPK282/0 cl4 parasites displays a trisomy for this chromosome when exposed to high SbIII concentrations^12^, suggesting that every chromosome has the potential to became polysomic depending on environmental pressures. The fact that, in our data, the karyotypes that dominate the population are similar to each other while karyotypes with several somy changes happen at low frequencies also suggests that selective pressure play a role in somy variability.

### Hypothesis on the evolution of mosaic aneuploidy in the clonal BPK282 population

By performing a pairwise comparison of the identified karyotypes in BPK282, we revealed a network structure where most karyotypes are linked to each other by single somy changes. This network allows proposing a hypothesis for the evolution of mosaicism during the 20 passages that followed cellular cloning. Kar1, the most frequent in the sequenced cells, is also the one displaying the highest number of karyotypes directly linked to it (**Figure 4**). Kar1 shows the same aneuploidy pattern as the ‘average’ karyotype assessed by BGS in the uncloned BPK282/0 line^11^, suggesting that this was also the dominant karyotype in the parental population. The high frequency of this karyotype in the parental population naturally increases the chance of it being randomly picked when starting a clonal population. Altogether, these data suggest that kar1 was the potentially founder karyotype of the BPK282 clonal population derived from this uncloned BPK282/0 line. Kar1 is likely well adapted to *in vitro* condition and it further spread during clonal propagation. According to the network, kar1 would have generated-through changes in somy of single chromosomes-a series of ‘primary’ derived karyotypes, (i) some diverging early and being quite successful like kar2-4 and (ii) others having diverged later and/or being less fit like the 19 minor karyotypes present around kar1. In this sense, there would be secondary, tertiary waves of karyotype diversification, like kar5 emerging from kar2 and kar15 emerging from kar5. Time lapse single cell sequencing would be required to test this hypothesis.

### Drug pressure and aneuploidy

In cancer cells and in human pathogenic yeast, genetic diversity created by mosaic aneuploidy has been linked to drug resistance^34,35^. In *Leishmania*, several *in vitro* studies report whole chromosome somy changes as one of the first genetic changes in populations under drug pressure^12,13,27,36,37^. However, it is still unknown if such variation in population average aneuploidy is a reflect of a positive selection of a sub-population displaying a pre-existing karyotype or is acquired *de novo* as a response to the pressure. The fact that the BPK282 population was previously used in *in vitro* drug selection experiments with at least 3 drugs^12,13,27^ allowed us to revisit the data of these experiments and compare the observed populational aneuploidy pattern changes with single karyotypes identified by SCGS in the population in order to address this question. In this context, we observed that the first contact of the parasites with the drugs lead to the emergence of new ‘populational karyotypes’ that are similar to karyotypes which were found in single-cells in the SCGS data (**Figure 5**). Thus, it seems that early stages of aneuploidy changes are caused by selection of pre-existing karyotypes. However, further exposure to higher concentrations led to somy changes that does not correspond to any karyotype in our data. It is possible that these changes reflect selection of other, rarer karyotypes that occur at frequencies lower than the detection limit of the droplet-based method applied here. Nonetheless, since *Leishmania* is able to constitutively generate mosaic aneuploidy^38^, it is also possible that new karyotypes are generated *de novo* as a response to the increasing challenge imposed by higher drugs concentration. Clonal lineages tracking methods, such as DNA barcodes^39^, should allow further insights in this matter.

## Conclusions

In summary, this work represents the first description of complete karyotypes of individual *Leishmania* parasites and open an unprecedent technological milestone to study MA in trypanosomatids. Future work should focus on the emergence and evolution of MA during clonal expansion by sequencing cells during very early stages of clonal evolution at different timepoints, as well as following the dynamics of karyotype changes during different stages of adaptation to drug pressure. Combining SCGS with single-cell transcriptomics could also allow to understand better the impact of gene dosage imbalance on transcription with a single cell resolution. Thus, high throughput single-cell sequencing methods represent a remarkable tool to understand key aspects of *Leishmania* biology and adaptability.

## Supplementary Material

**Figure S1.**
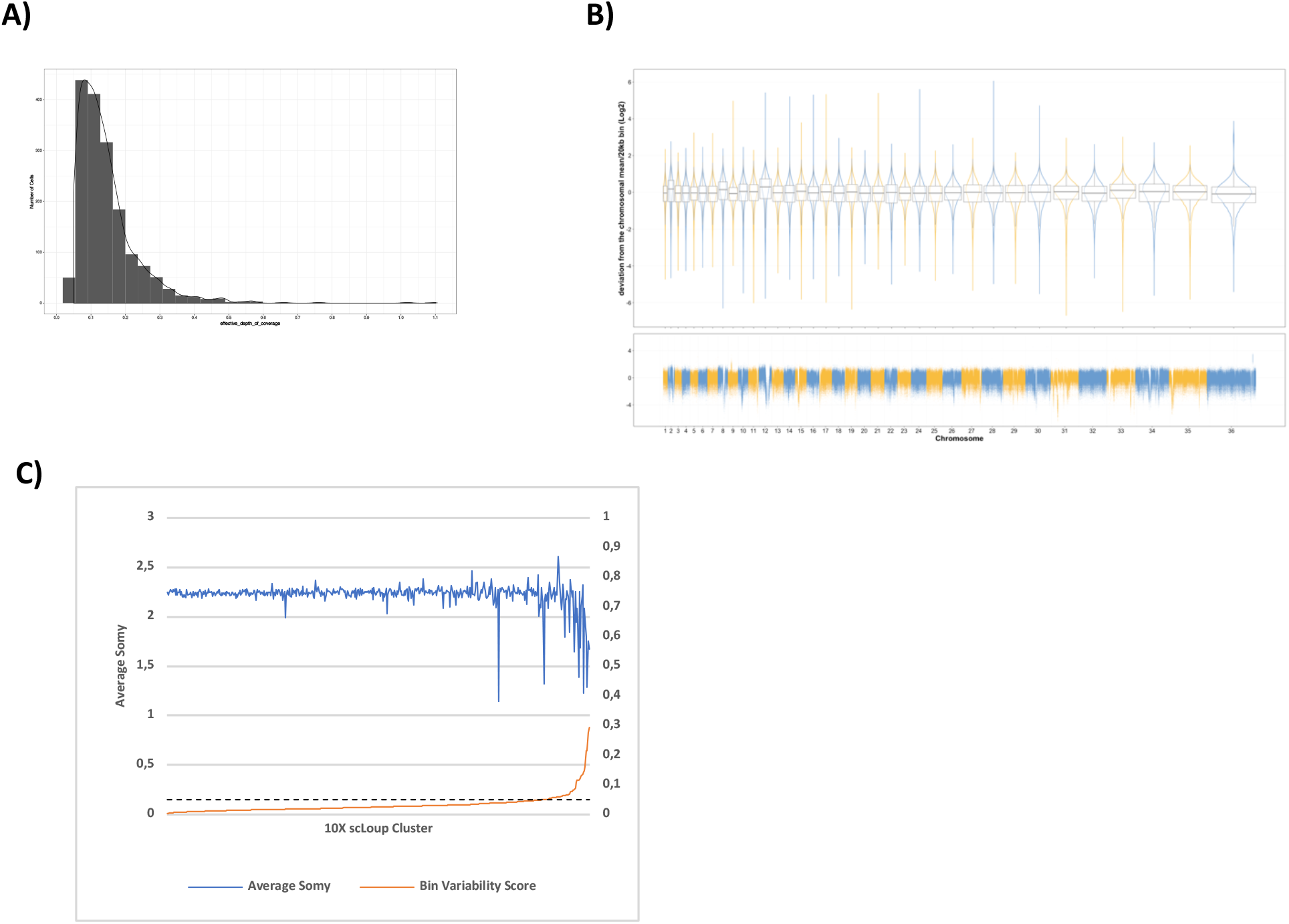
**A)** Distribution of the effective coverage depth per cell. **B)** Violin plot showing the read depth deviation from the mean of intrachromosomal 20kb bins. For each cell, the mean depth/20kb of each chromosome was calculated and the ration between the depth of each intrachromosomal 20kb bin and the average was log2 transformed. Values close to 0 represent bins with depth similar to the chromosomal average. A box plot indicates the median (line in the center), the upper and lower quartiles (boxes) and the maximum and minimum of each chromosome (lines). A Manhattan plot on the bottom shows individual deviation values. **C)** Average somy and bin variability score of all 512 clusters. The black dashed line represents the threshold set to eliminate clusters with high bin-to-bin variability.

**Figure S2.**
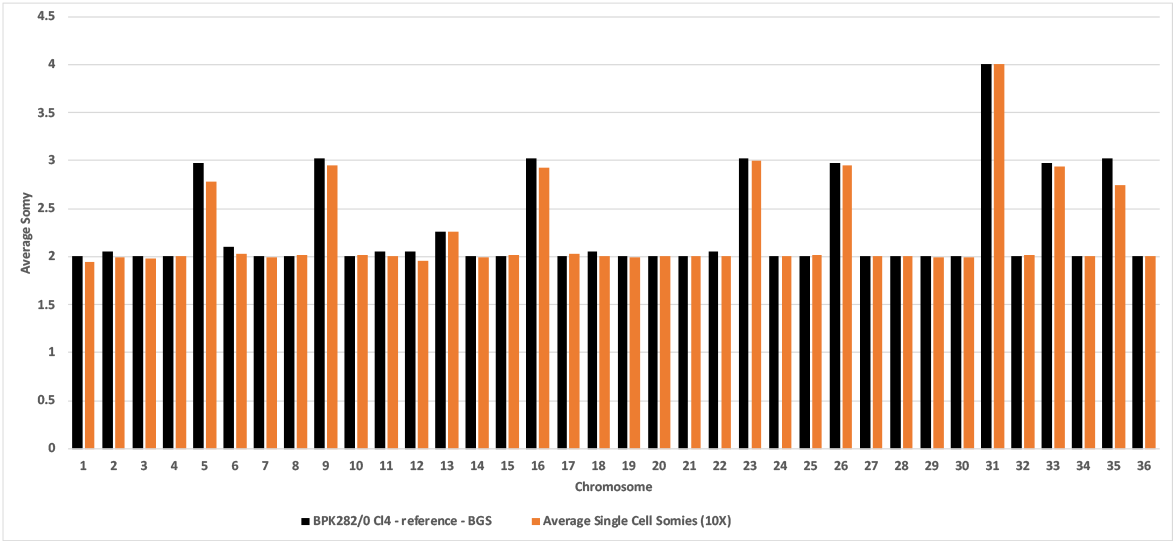
Comparison of the average combined somy values of all 1560 BPK282/0 cl4 promastigotes assessed by SCGS (orange bars) with the average somies of a population of the reference BPK282 strain estimated by BGS (black bars).

**Figure S3.**
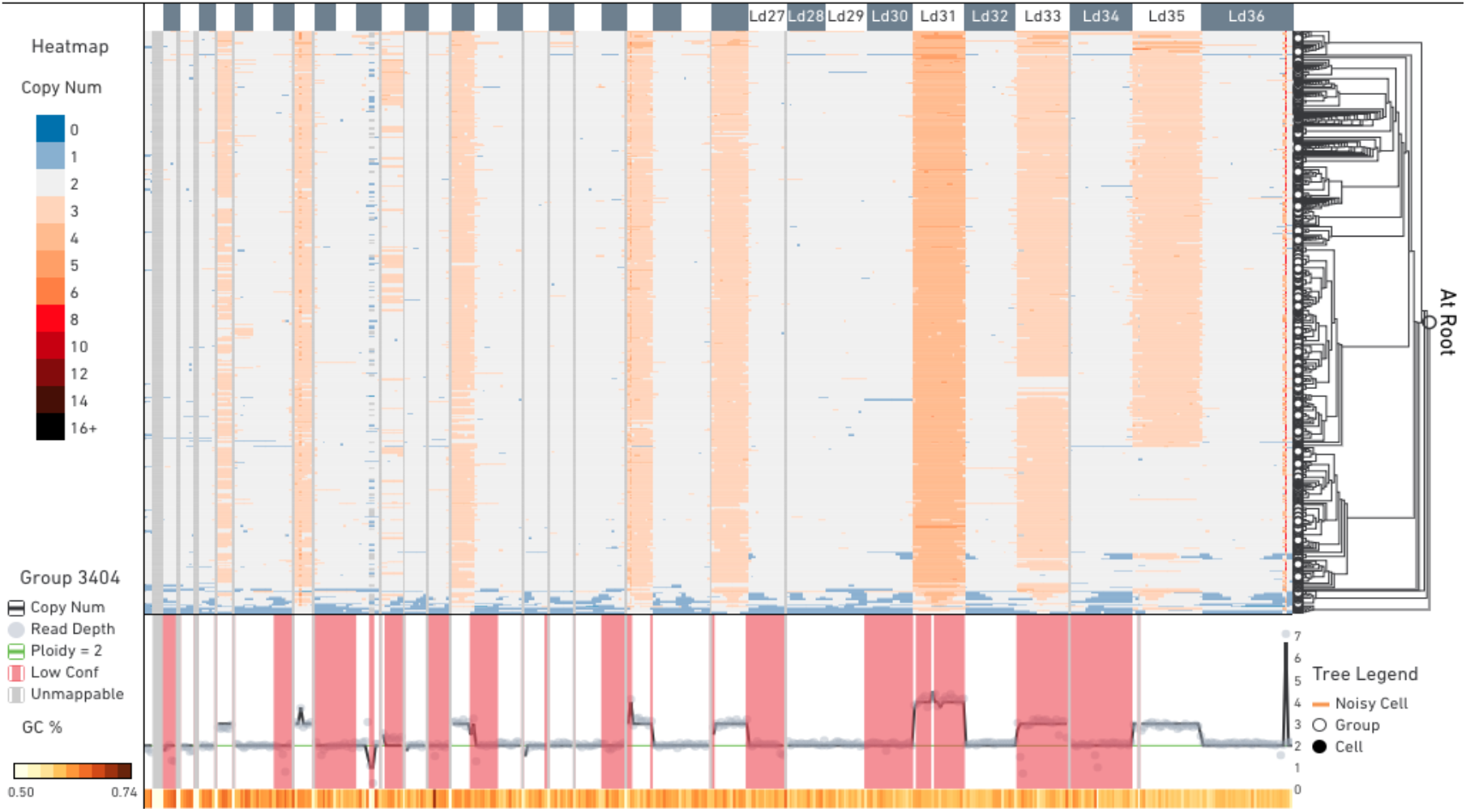
CNV values calling by 10X Cell Ranger DNA pipeline visualized in 10X Loupe scDNA Browser tool. The 1703 cells are arranged in 512 clusters (horizontal lines). Bars on the top indicate the position of each chromosome, while lines and grey dots in the bottom represent the CNVs and reads/megabase, respectively, of all clusters combined. High copy number regions in chromosome 23 and 36 represent, respectively, the H-locus and the M-locus.

**Figure S4.**
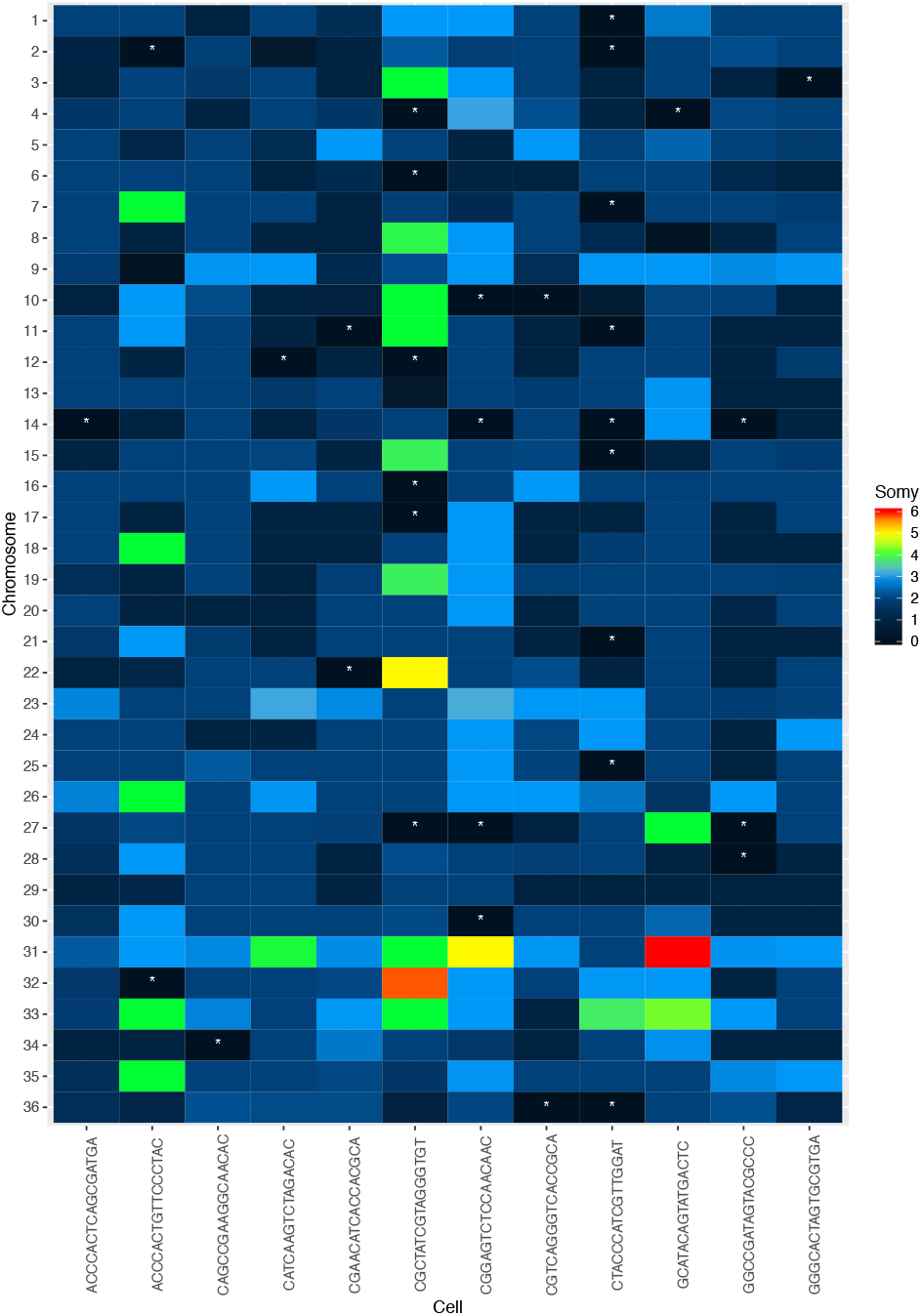
Heatmap showing the estimated somy of all cells where at least one chromosome was missing, including cells removed from by data filtering. Nullisomic chromosomes are marked with a white asterisk.

## Notes

### Competing Interest Statement

The authors have declared no competing interest.

### Summary of Updates

Added statement on page 2.

